# Allosteric Regulation of Aptamer Affinity through Mechano-Chemical Coupling

**DOI:** 10.1101/2022.07.28.501929

**Authors:** Hao Qu, Manyi Zheng, Qihui Ma, Lu Wang, Yu Mao, Michael Eisenstein, Hyongsok Tom Soh, Lei Zheng

## Abstract

The capacity to precisely modulate aptamer affinity is important for a wide variety of applications. However, most such engineering strategies entail laborious trial-and-error testing or require prior knowledge of an aptamer’s structure and ligand-binding domain. We describe here a simple and generalizable strategy for allosteric modulation of aptamer affinity by employing a double-stranded molecular clamp that destabilizes aptamer secondary structure through mechanical tension. We demonstrate the effectiveness of the approach with a thrombin-binding aptamer and show that we can alter its affinity by as much as 65-fold. We also show that this modulation can be rendered reversible by introducing a restriction enzyme cleavage site into the molecular clamp domain and describe a design strategy for achieving even more finely-tuned affinity modulation. This strategy requires no prior knowledge of the aptamer’s structure and binding mechanism and should thus be generalizable across aptamers.

## Introduction

Affinity reagents such as antibodies and aptamers are essential tools in both basic and applied research for a wide variety of functional materials and molecular diagnostics systems.^[1]^ The operating parameters of these molecules are inherently constrained by the thermodynamic and kinetic binding properties of the affinity reagent being employed. However, the usefulness of such reagents can be further extended by introducing mechanisms that make it possible to tune their affinity—examples of such applications include biosensors with extended dynamic ranges,^[2]^ accurate tracking for molecular imaging,^[3]^ precise loading and delivery of drugs,^[4]^ fine control of nanodevices,^[5]^ and regulation of gene expression.^[6]^ The binding properties of affinity reagents are inherent to the nucleic acid or amino acid sequence and structure generated during the screening and selection process, and further engineering is therefore required to manipulate that structure and thereby achieve the desired effect on ligand affinity. In general, this is a much more straightforward process for aptamers relative to antibodies, because their structure can be readily engineered based on predictable base-pairing principles. Furthermore, researchers have identified a number of well-known and -defined structural motifs, such as hairpins and G-quadruplexes, which can be incorporated into an aptamer-based system in order to alter its folding and binding properties in a semi-predictable fashion.

Affinity modulation is usually achieved through direct mutations in aptamer sequences, followed by extensive screening and validation procedures to locate variants with appropriately altered affinity.^[7]^ However, this mutagenesis-based approach is time-consuming and inefficient, due to its trial-and-error nature and because it is difficult to ensure that a given mutation will not only meaningfully alter affinity, but also achieve this by modifying the conformational equilibrium rather than the chemical interaction between the reagent and the analyte.^[8]^ As an alternative, one can rationally design aptamer-based molecular switch constructs with modulated affinity, which transition between a binding-ready state and a binding-inactive stem-loop structure.^[2a, 9]^ Effective affinity can also be controlled through the use of split aptamer constructs, in which affinity is modulated by the purely entropic change associated with the length of the intramolecular linker joining the two fragments.^[10]^ Alternatively, aptamer affinity can be modulated by adding a triple helix structure to the two termini of an aptamer, thereby trapping them in a fixed position and limiting the aptamer’s flexibility.^[11]^ However, these design strategies typically require precise knowledge of both the structure of the folded aptamer as well as the basis of its interaction with the ligand, limiting their general applicability. Several groups have described strategies in which an aptamer is reversibly stabilized in a non-binding conformation by using a complementary strand that hybridizes to a portion of the aptamer as an allosteric inhibitor.^[2b, 2c, 12]^ One important limitation of this allosteric approach is that the inhibitor strands typically interact with the active binding domain in direct binding competition with the ligand. Unfortunately, this approach is challenging to implement for applications in which the goal is to partially down-tune aptamer affinity by a defined amount rather than inhibit aptamer binding entirely.

In this work, we describe a simple and effective strategy for reversibly fine-tuning aptamer affinity by adding an allosteric ‘tuning clamp domain’ that acts at a distance from the binding site. Specifically, we use a molecular clamp design that perturbs aptamer folding through mechanical extension, forcing it into a state in which its binding competency is considerably reduced. This molecular clamp structure self-assembles through the hybridization of an allosteric inhibitor DNA strand with two clamp domain sequences flanking the aptamer. We also describe simple design strategies that make it possible to precisely and predictably tune the affinity of a given aptamer by a defined amount, along with a restriction enzyme-based strategy for releasing the tension of the molecular clamp and restoring normal aptamer affinity. Previous work has demonstrated the nanomechanical control of enzymatic activity,^[13]^ but to our knowledge, this represents the first attempt to modulate aptamer affinity in such a fashion. As a proof of concept, we have modulated the affinity of HD22, an aptamer for human α-thrombin,^[14]^ by ~65-fold, and show that the mechanically-inhibited HD22 aptamer can be reactivated and restored to its initial affinity via endonuclease digestion. Given that no prior knowledge of the aptamer structure or binding domain is required, we believe this strategy should be generally applicable across a range of aptamers.

## Results and Discussion

### Design principles for the tuning clamp domain system

Our clamp design consists of two strands (**Fig. 1a**). The first strand is composed of the aptamer flanked by ‘clamp domain’ sequences at both ends. These flanking sequences are designed to hybridize with a complementary allosteric inhibitor strand, forming a D-shaped, mechanically-constrained ‘stressed aptamer molecule’ (SAM) (**Fig. 1b**). The SAM can be viewed as two mechanically coupled, nonlinear springs: a double-stranded molecular clamp and the aptamer. These are constrained at the same end-to-end distance (EED), which reflects the distance between the two ends of the SAM and thus describes the degree of deformation for both components. The aptamer is driven into the unfolded state by the hybridization of the allosteric inhibitor strand, which forms the molecular clamp structure and thereby shifts the aptamer population from a folded state in which it can bind to the target to an unfolded state with greatly reduced affinity (**Fig. 1b**). By modulating the stiffness of the clamp, we anticipated that it should be possible to alter the proportion of the aptamer that exists in this unfolded state, enabling systematic regulation of aptamer affinity.

**Figure 1.**
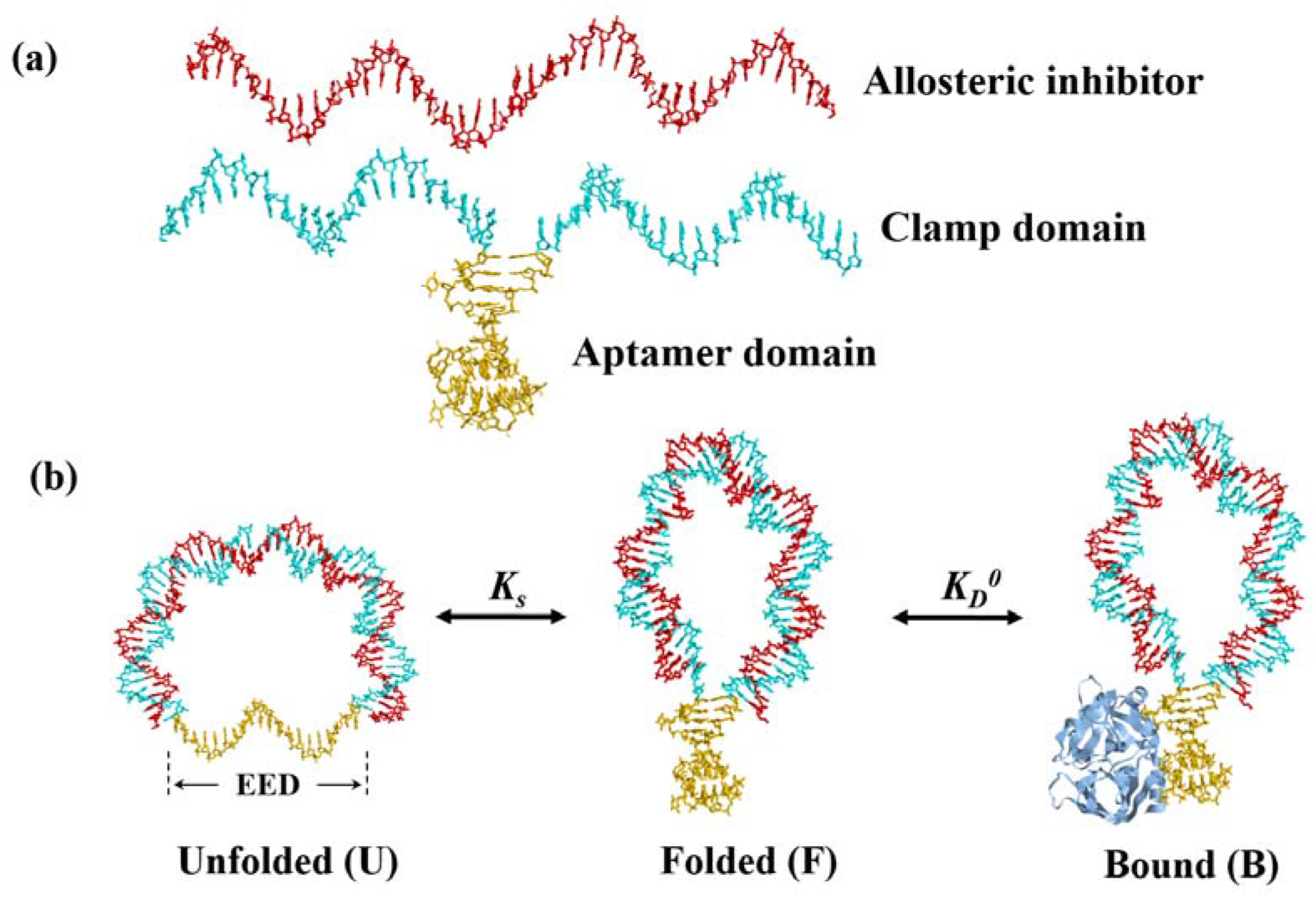
Schematic of allosteric control with the tuning clamp domain. (**a**) An aptamer sequence (yellow) is flanked by two clamp domain sequences (cyan). These domains are designed to hybridize with a complementary allosteric inhibitor strand (red). (**b**) When fully assembled, the two strands form a stressed aptamer molecule (SAM). This forces the aptamer into an unfolded state (U), shifting the equilibrium away from the folded, binding-competent (F) and target-bound (B) states.

The influence of the unfolded state on overall aptamer affinity can be analytically determined based on the population shift model.^[2b]^ Briefly, the dissociation constant (*k*_*D*_) of SAMs for the aptamer target can be described by the following equation:

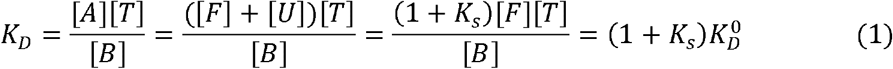

where [A], [T], [B] are the respective concentrations of aptamer, target, and aptamer-target complex, [F] and [U] are the concentration of the folded and unfolded states, *K*_*D*_^0^ is the initial dissociation constant of the parent aptamer, and *K*_*S*_ is the interconversion rate between the folded and unfolded states. This final term is described by:

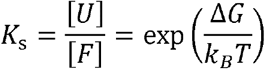

Thus, the, *K*_*D*_ of the SAM can be described as follows:

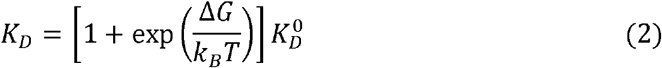

in which is *k*_*D*_ the Boltzmann constant, *T* is the temperature, and Δ*G* is the free energy difference between the folded and unfolded states. Based on Eq. (2), it is clear that one can control aptamer affinity by tuning the Δ*G* value. To determine Δ*G* quantitatively for the SAM, one can calculate the total energy profile of the SAM using the bending energy model for the molecular clamp and the stretching energy model for the aptamer (see Methods). We demonstrated this with a series of modeling experiments based on SAMs incorporating the thrombin-binding aptamer HD22.^[14]^ When we plotted the total elastic energy (E_tot_) of the system as a function of EED (**Fig. 2a**), we identified two local minima that correspond to the folded and unfolded states of the aptamer. The difference between these two minima yields the Δ*G* value. A stiffer molecular clamp will exert greater force on the aptamer and will therefore make the unfolded state more energetically favorable. We therefore expect that the balance between the folded and unfolded states can be shifted by changing the stiffness of the molecular clamp, which can in turn be adjusted by altering its length (*N*_*d*_), as shown in our previous work.^[15]^ To analyze the influence of the clamp length, we plotted the total energy profiles of SAMs with molecular clamps ranging in length from 16 to 24 bp (**Fig. 2b)**. The simulated constructs are denoted as SAM16-16, SAM18-18, SAM20-20, SAM22-22, and SAM24-24, where the first number refers to the length of the tuning clamp domain and the latter number refers to the length of the allosteric inhibitor strand. The energy minimum for the folded state was barely changed, while that for the unfolded state was dramatically shifted, suggesting that Δ*G* can be readily regulated by varying *N*_*d*_.

**Figure 2.**
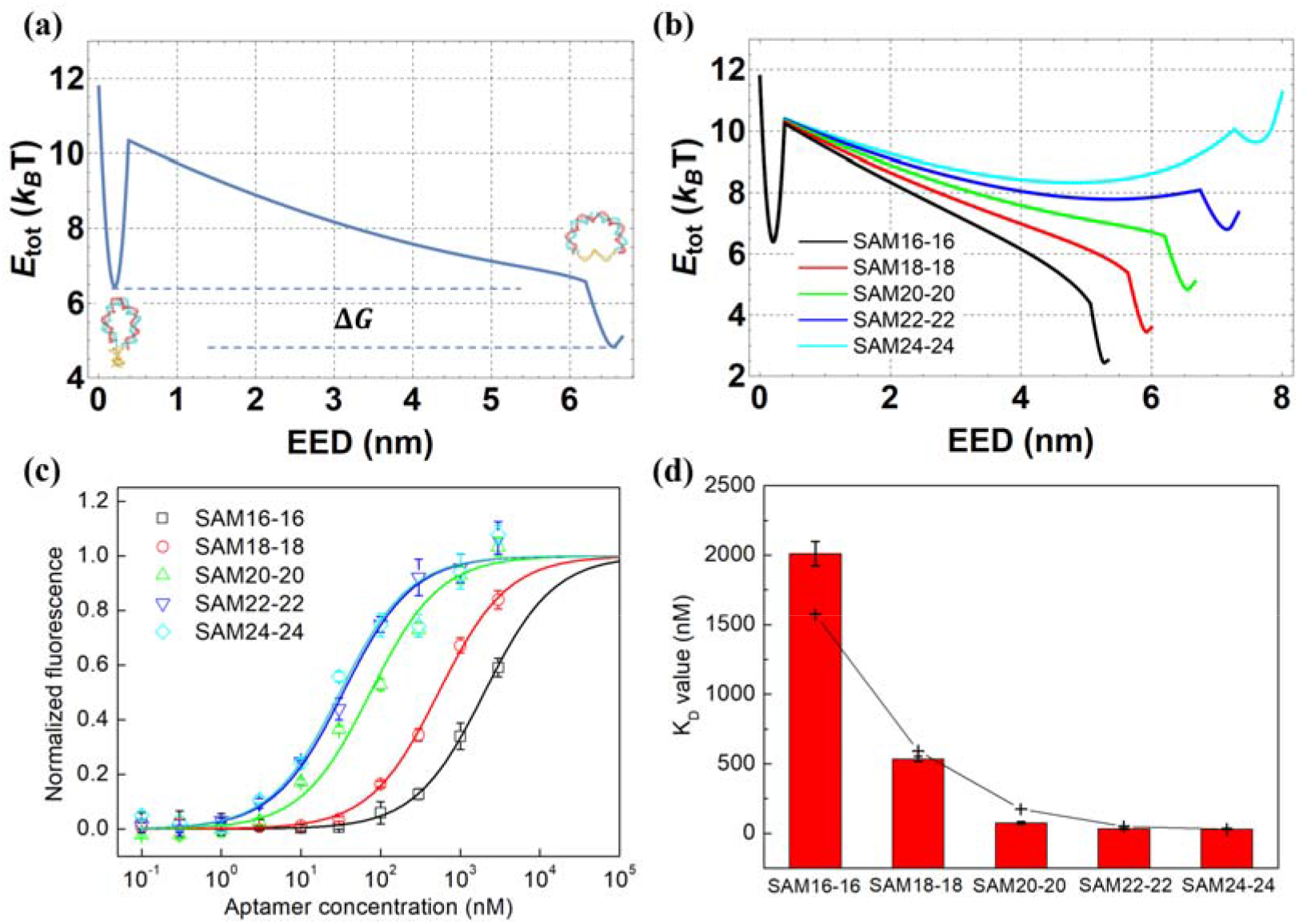
Regulating aptamer affinity by tuning the mechanical properties of the molecular clamp. (**a**) Predicted total energy (E_tot_) profile vs EED for various SAM constructs, using the bending energy model for the molecular clamp and the stretching energy model for the aptamer. The two energy minima correspond to the folded (left) and unfolded (right) states of the aptamer. (**b**) Predicted total energy profiles for SAMs with molecular clamp length (*N*_*d*_) ranging from 16 to 24 bp. (**c**) Binding curve modulation using molecular clamps of varying lengths. (**d**) The aptamer *K*_*D*_ values extracted from the binding curves in (**c**). The black crosses are the predicted *K*_*D*_ value based on Eq. (2), with the free energy difference *ΔG* extracted from the model shown in (**b**).

### Experimental validation of molecular clamp-based tuning of aptamer affinity

To verify these modeling-based predictions, we synthesized a series of HD22-based aptamer strands with tuning clamp domains of varying length (16, 18, 20, 22, and 24 nt). These were then hybridized with fully complementary allosteric inhibitor strands to produce SAMs with molecular clamps of *N*_*d*_ ranging from 16–24 bp. We measured the apparent *K*_*D*_ using a standard fluorescence binding assay, in which we incubated thrombin-coated beads with various concentrations of fluorescently labeled SAMs and determined the target-bound fraction with a microplate reader (see Methods). The results of this assay showed that SAMs with lower-stiffness molecular clamps (*N*_*d*_ = 24, 22, and 20) did not meaningfully alter the aptamer binding curve, whereas stiffer molecular clamps (*N*_*d*_ = 18 and 16) dramatically shifted the curve to the right (**Fig. 2c**). We derived the *K*_*D*_ values for these various constructs by fitting the binding curve to the Langmuir isotherm (**Fig. 2d**), and determined that the affinity of the SAM16-16 construct (2,010.88 ± 87.33 nM) was reduced by nearly 65-fold relative to the SAM24-24 construct (30.95 ± 5.67 nM). These results confirmed that one can achieve effective modulation of aptamer affinity through this molecular clamp design.

We then calculated the total elastic energy profiles for these SAMs with an identical set of parameters: bending modulus *B* = 50 k_B_T or 200 pN × nm^2^, critical torque τ_c_ = 28 pN × nm for the dsDNA bending energy, persistence length *l*_*p*_ = 1.12 nm, and aptamer folding energy Δ*G*_fold_ = 4.34 k_B_T (as determined previously).^[16]^ The Δ*G* values were extracted from **Fig. 2b**, and the predicted *K*_*D*_ values were calculated using Eq. (2) as plotted in **Fig. 2d**. Remarkably, this single set of parameters showed an excellent fit for the experimental *K*_*D*_ values of all the tested SAMs.

### Releasing the mechanical tension in SAMs to restore normal aptamer binding

The above experiments strongly indicate that aptamer affinity can be directly modulated by introducing changes in mechanical tension of the molecular clamp, and we subsequently validated this in series of control experiments. Every SAM features a gap in the middle of the molecular clamp structure, where the flanking arms of the aptamer strand meet (*i*.*e*., the space between the two cyan domains in **Fig. 1b**). In principle, widening this gap should release the mechanical tension applied to the aptamer (**Fig. 3a**), since most of the elastic energy in the molecular clamp is stored in the middle of the double-stranded DNA domain.^[17]^ We tested this by combining an aptamer strand with a 16-nt tuning clamp domain with various allosteric inhibitor strands that were designed to extend the width of this gap. Our experiments confirmed that as the gap increases in length from 0 to 3 nt, the aptamer *K*_*D*_ dropped back to *K*_*D*_^0^ (22.38 ± 3.48 nM; **Fig. 3b, c**), clearly indicating that allosteric control of aptamer affinity in the context of the SAM was being achieved through the strain exerted by the molecular clamp.

**Figure 3.**
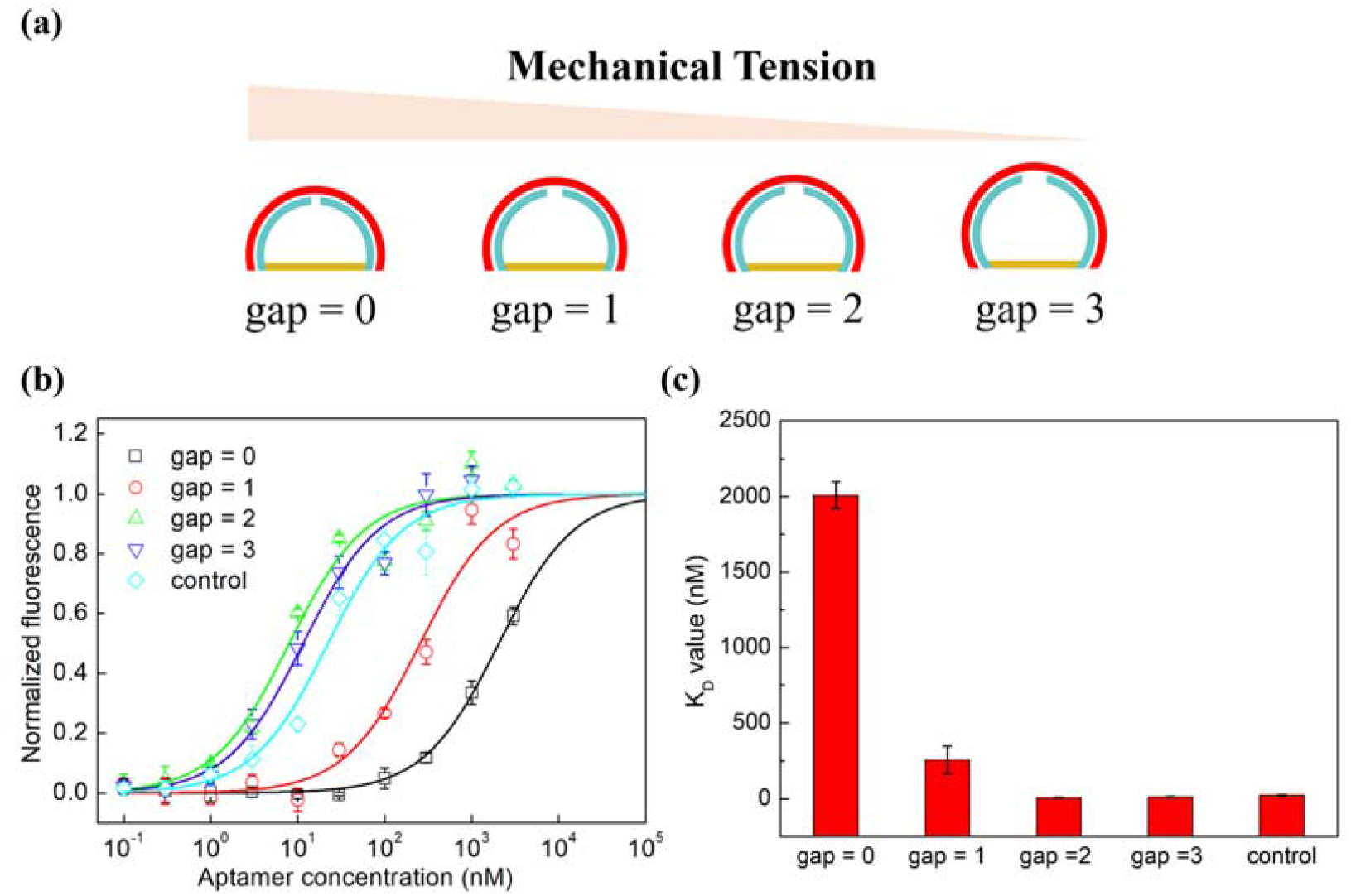
Mechanical tension is the key to molecular clamp-mediated regulation of aptamer binding. (**a**) Schematic showing how the size of how single-stranded gaps in the molecular clamp affect mechanical tension **(b)** The thrombin binding curve can be modulated by increasing the single-stranded gap at the center of the molecular clamp from 0 to 3 nt. Control is the aptamer strand alone, without the allosteric inhibitor strand. (**c**) *K*_*D*_ values extracted from the binding curves in (**b**), ranging from 2,010.88 ± 87.33 nM for a 0 nt gap to 11.58 ± 2.87 nM for a 3 nt gap.

We also identified a mechanism to efficiently alleviate this mechanically-induced allosteric regulation, thereby restoring the baseline affinity of the aptamer. To achieve this, we incorporated an EcoRI restriction endonuclease site into the molecular clamp region of construct SAM16-16. After hybridization, we measured a *K*_*D*_ value of 1,692.55 ± 184.03 nM, demonstrating strong inhibition of the aptamer. But after digestion with EcoRI, the aptamer was restored to near-baseline affinity, with a *K*_*D*_ of 56.33 ± 5.48 nM (**Fig. 4**), indicating successful disassembly of the molecular clamp.

**Figure 4.**
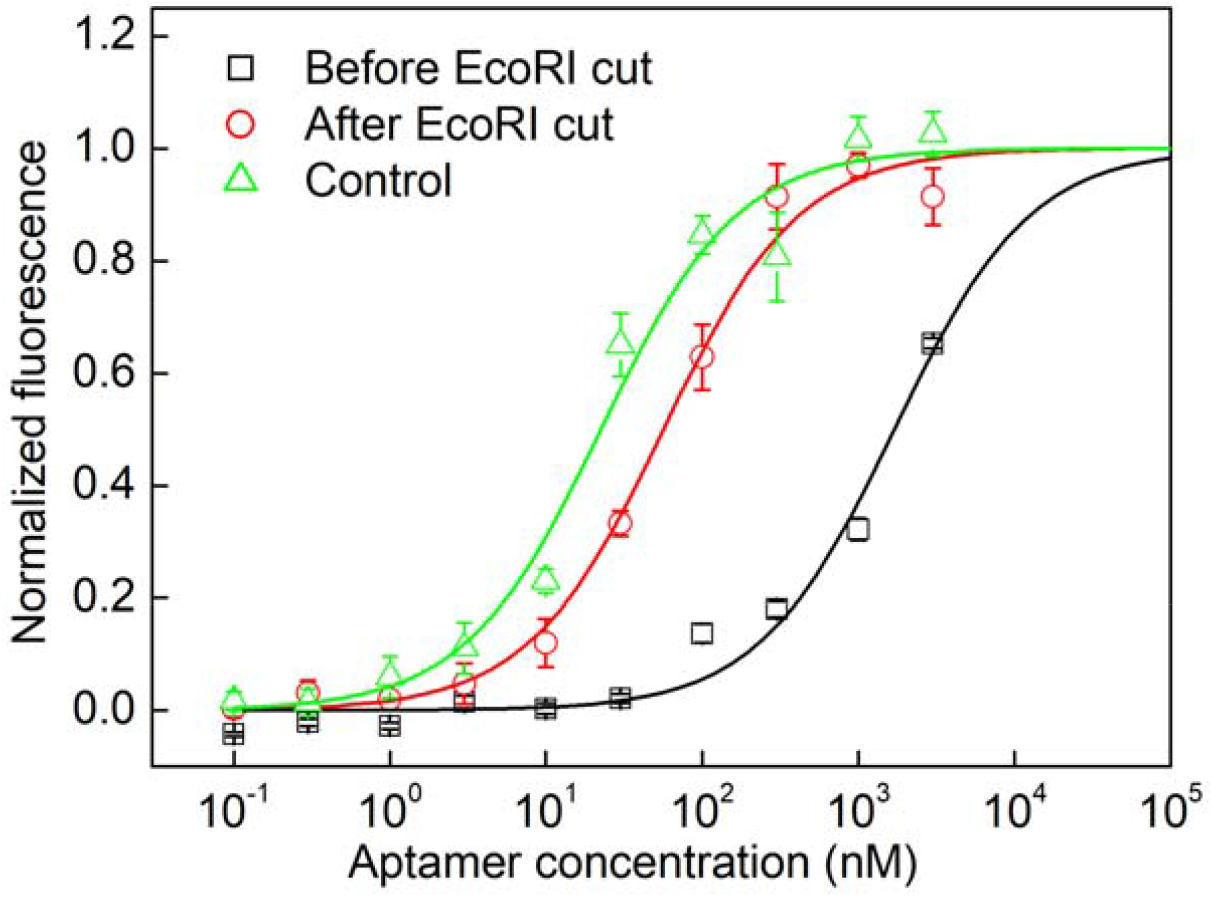
Binding curves show restoration of aptamer affinity after EcoRI digestion of the molecular clamp. The *K*_*D*_ value shifted from 1,692.55 ± 184.03 nM to 56.33 ± 5.48 after targeted cleavage of the molecular clamp domain. The control curve is derived from the aptamer strand in the absence of the allosteric inhibitor strand.

### Fine-tuning of aptamer affinity using single-stranded ‘hinges’

It is worth noting that according to Eq. (2), when Δ*G* < 0, *K*_*D*_ ≈ *K*_D_^0^, but when Δ*G* > 0, *K*_*D*_ increases exponentially with Δ*G*, and we also observed this in our experimental results (**Fig. 2d**). These results indicate that this simple molecular clamp system is not necessarily suitable for precisely fine-tuning an aptamer’s target-binding affinity on its own. We subsequently devised a mechanism for achieving such fine calibration, wherein we introduced single-stranded ‘joints’ between the molecular clamp and the aptamer that partially release the internal energy of the SAM. We achieved this by using shorter allosteric inhibitor strands that are only complementary to a partial stretch of the aptamer strand’s tuning clamp domain. For example, an aptamer strand with a clamp domain of 22 nt might be hybridized with a 16-nt allosteric inhibitor strand (SAM22-16), yielding a SAM with a molecular clamp of *N*_*d*_ = 16 bp flanked by a 3-nt single-stranded joint on either side. We expected that for any given length of the clamp domain, the existence of flanking single-stranded joints would act as a hinge that releases some of the mechanical tension applied to the aptamer and thereby reduces the effect of the molecular clamp on the *K*_*D*_ value. As these single-stranded joints increase in length from 1 to 3 nt, we would expect this to further soften the flanking hinge and thereby enhance this dampening effect on the molecular clamp (**Fig. 5a**).

**Figure 5.**
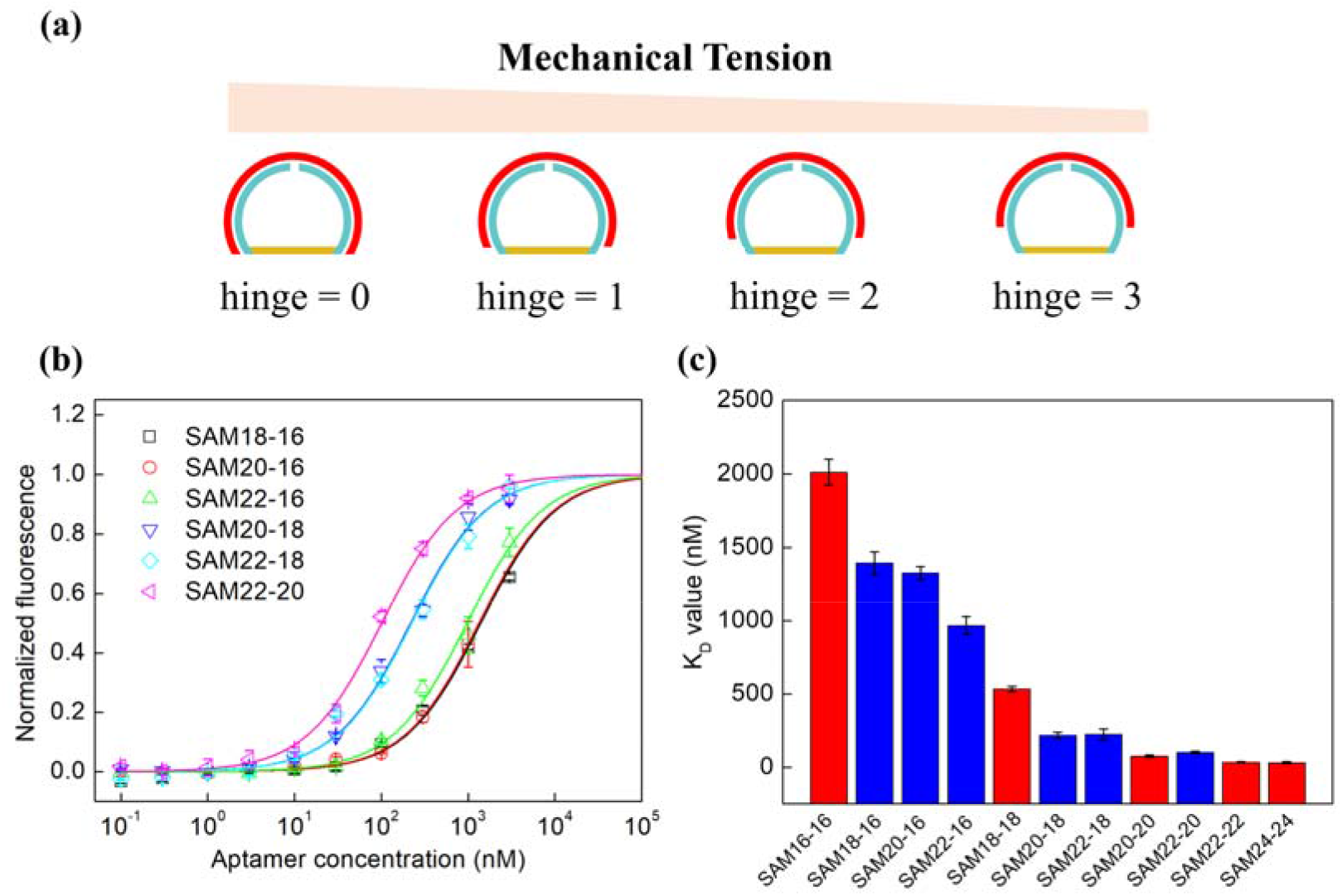
Fine-tuning aptamer affinity by adding ‘hinges’ adjacent to the molecular clamp domain. (**a**) Schematic showing how single-stranded hinges of increasing length affect SAM mechanical tension. (**b**) Binding curves for a variety of SAMs containing single-stranded hinges at either end of the tuning clamp domain. (**c**) Affinity measurements for fully complementary SAMs (red) and constructs that contain flanking hinges (blue).

In order to test this directly, we constructed and generated binding curves for a number of SAMs with varying lengths of molecular clamp and single-stranded joint regions: SAM18-16, SAM20-16, SAM20-18, SAM22-16, SAM22-18, and SAM22-20 (**Fig. 5b**). The affinity measurements for these various SAMs (blue bars in **Fig. 5c**) confirmed our initial predictions, and we consistently observed *K*_*D*_ values that were lower than those of the equivalent SAMs containing no single-stranded joints (red bars in **Fig. 5c**). The affinity achieved with these SAMs with single-stranded joints effectively filled the gaps in **Fig. 2d**, clearly demonstrating that this fine-tuning strategy allows us to better access the full range of possible affinities that can be achieved with our SAMs.

## Conclusion

We present a strategy for aptamer affinity regulation based on an intuitive design that requires only the addition of tuning clamp domain sequences at both ends of the aptamer and the synthesis of a complementary allosteric inhibitor strand. The allosteric inhibitor strand forms a mechanical module that is physically distinct from the binding domain of the aptamer, avoiding direct competition with the ligand, thereby achieving affinity regulation in a straightforward and predictable fashion. The allosteric inhibition of aptamer binding can be rapidly alleviated in a controlled fashion by incorporating a restriction enzyme cleavage site within the tuning clamp domain sequence. Finally, we have demonstrated that we can achieve even finer control over the final affinity of the aptamer by introducing single-stranded ‘hinge’ domains that partially alleviate the mechanical tension from the molecular clamp. Importantly, this approach does not require any *a priori* knowledge of the aptamer’s structure or thermodynamic parameters, and we believe it should be broadly generalizable across aptamers.

This approach is not without limitations. The affinity tuning range is relatively small due to the intrinsic gap in the molecular clamp,^[17]^ although repair of this gap by cyclizing the aptamer strand should further broaden this range. Since circular aptamers have been reported to possess improved biostability and affinity,^[18]^ such constructs could prove valuable for the controlled delivery and release of drugs.^[19]^ Another limitation is that aptamer affinity can only be down-tuned using this molecular clamp system. Strategies that restrict aptamer flexibility through terminal fixation^[11]^ or by manipulate the trade-off between aptamer affinity and cooperativity^[20]^ could offer solutions for allosterically activating aptamer affinity, and this will be an interesting avenue for future investigations.

## Acknowledgements

This study was supported by the National Natural Science Foundation of China (32172294), and the Key Science & Technology Specific Projects of Anhui Province (202003a06020017). This work was also supported by the Leona M & Harry B. Helmsley Charitable Trust, (HTS). We thank Prof. Feng Wang for assistance with the microplate reader.

