## Supporting Information for "Allosteric Regulation of Aptamer Affinity through Mechano-Chemical Coupling"

**through Mechanochemical Coupling**

**Methods and Materials**

### *Chemicals and reagents*

Human α-thrombin (Factor IIa) was purchased from Yeasen Biotechnology (Shanghai, China). Dynabeads (M-270 carboxylic acid) were obtained from Thermo Fisher. N-hydroxy succinimide (NHS), 1-(3-dimethylaminopropyl)-3-ethylcarbodiimide hydrochloride (EDC), phosphate-buffered saline (PBS), Tween 20, and EcoRI restriction enzyme were purchased from Sangon Biotech (Shanghai, China). All other chemicals and reagents were purchased from Sinopharm (Shanghai, China).

### *Sample preparation for SAMs*

All oligonucleotide sequences (listed in **Table S1**) were synthesized by Sangon Biotech with HPLC purification. We confirmed hybridization of dsDNA regions by checking sequence thermodynamic properties with the DINAMelt web server ^[^^1^^]^. Each aptamer strand was mixed with equimolar amounts of allosteric inhibitor strand and diluted to a final concentration of 2 μM in 200 μL hybridizing buffer (10 mM Tris, 5 mM MgCl_2_, 100 mM NaCl, pH 7.9). The mixture was then heated to 95 ℃ for 10 min and annealed at room temperature overnight to ensure the proper conformational structures in the stressed DNA molecules.

### *Determination of dissociation constants (K_D_)*

*K_D_* values were measured using a standard fluorescence binding assay. Thrombin-coated Dynabeads were prepared following the manufacturer’s protocol for two-step coupling using EDC and NHS. Next, 7 × 10^5^ of the thrombin-coated beads were washed three times in PBSMT buffer (1 mM MgCl_2_, 100 mM NaCl, 4 mM KCl, 0.025% Tween 20) and then incubated with freshly prepared FAM-labelled SAMs at final concentrations ranging from 0.1 nM to 3 μM for 40 min at room temperature with gentle rotation. The unbound SAMs were washed away with three washes of PBS buffer. The bound fraction of SAMs was determined using a microplate reader (Tecan Infinite M Plex) with excitation at 485 nm and emission at 525 nm. The binding curve of the SAMs (fluorescence intensity *vs* concentration) was fitted using the nonlinear equation $\text{y = B}\left（ \text{1 + }{\text{K}_{\text{D}}}/\text{x} \right）\text{ + C}$ to extract *K_D_* values.

### *Digestion of the molecular clamp by EcoRI*

SAM16-16 (**Table S1**) was prepared following the above protocol for SAM assembly. 200 μL of 2 μM SAM16-16 was divided evenly into two tubes. One aliquot was mixed with 6.5 μL EcoRI restriction enzyme (5 kU) and the other was mixed with 6.5 μL 1× PBSMT as control. Both samples were incubated at 37 ℃ for 16 h to ensure full digestion of the molecular clamp by EcoRI.

### *Theoretical principle of the method*

The extension of the dsDNA molecular clamp creates an unfolded state for the aptamer. According to our previous studies^[^^2^^]^, the bending elastic energy of the molecular clamp can be analytically expressed by:

$$E_{d}\left( x \right)=\left\{ \begin{aligned} \tau_{c}\arccos\left( \frac{x}{2R} \right), 0<x<x_{c} \\ \frac{5B}{L}\frac{x_{0}-x}{2L}-k_{B}T\ln\left( \frac{2L-x}{2L-x_{0}} \right), x_{c}<x<x_{0} \end{aligned} \right. (S1)$$

where τ_c_ is the critical torque at which the kink develops; *x* the end-to-end distance (EED); $R=L(1-2\gamma^{2}/45)$ is the arc length, where $\gamma=L\tau_{c}/(2B)$, 2*L* is the contour length of the molecular clamp, and *B* is the bending modulus; $x_{0}=2L[1-k_{B}TL/(5B)]$ is the EED at zero force; and *x_c_* is the critical EED. The lower form of the equation corresponds to the smoothly bent solution, while the upper describes the kinked state. Therefore, the bending behavior – including the applicable force for using short dsDNA as molecular clamp – is comprehensively known. For example, according to Eq. S1, the bending elastic energy of dsDNA shows an increase with decreasing length (*N_d_*). Hence, the stretching state of the aptamer can be readily tuned by adjusting *N_d_*.

The stretching force applied to the aptamer in the context of the clamp initially leads to disruption and unfolding of its structure, with a corresponding free energy $\Delta G_{fold}$. After the aptamer is completely unfolded as single-stranded DNA without any internal structure, the free energy required for pulling is purely elastic, arising from the entropy change of the single-stranded DNA as a random coil ($\Delta G_{coil}$). The random coil state follows the Worm-like-Chain (WLC) model described by the Marko-Siggia expression ^[^^3^^]^:

$$\Delta G_{coil}=\int_{0}^{x} \frac{1}{l_{s}}\left[ \frac{3x}{N_{s}}+\frac{1}{4\left( 1-\frac{3x}{N_{s}} \right)^{2}}-\frac{1}{4} \right]dx (S2)$$

where *x* is the EED, *N_s_* the number of bases of single-stranded DNA, and *l_s_* the persistence length of single-stranded DNA. Therefore, the stretching energy for the aptamer can be modeled as the superposition of an energy well with a depth of $\Delta G_{fold}$ in the elastic energy of random coil $\Delta G_{coil}$, as described in Eq. S2. For the HD22 aptamer used in this work, we have previously found that $\Delta G_{fold}=4.34 k_{B}T$. Note that the detailed energy landscape of aptamer folding energy is not discussed in this paper.

**Table S1.** **DNA sequences used in this work**

|  | Aptamer strand (5’ - 3’) | Allosteric inhibitor strand (5’ - 3’) |
| --- | --- | --- |
| SAM24-24 | ACGTGAGAGCAGAGTCCGTGGTAGGGCAGGTTGGGGTGACT GACATACGACGA | [FAM]-CTGCTCTCACGT TCGTCGTATGTC |
| SAM22-22 | ACGTGAGAGCAAGTCCGTGGTAGGGCAGGTTGGGGTGACT ACATACGACGA | [FAM]-TGCTCTCACGT TCGTCGTATGT |
| SAM20-20 | ACGTGAGAGCAGTCCGTGGTAGGGCAGGTTGGGGTGACT CATACGACGA | [FAM]-GCTCTCACGT TCGTCGTATG |
| SAM18-18 | ACGTGAGAGAGTCCGTGGTAGGGCAGGTTGGGGTGACT ATACGACGA | [FAM]-CTCTCACGTTCG TCGTAT |
| SAM16-16 | ACGTGAGAAGTCCGTGGTAGGGCAGGTTGGGGTGACT TACGACGA | [FAM]-TCTCACGTTCG TCGTA |
| SAM18-16 | ACGTGAGAGAGTCCGTGGTAGGGCAGGTTGGGGTGACT ATACGACGA | [FAM]-TCTCACGTTCG TCGTA |
| SAM20-16 | ACGTGAGAGCAGTCCGTGGTAGGGCAGGTTGGGGTGACT CATACGACGA | [FAM]-TCTCACGTTCG TCGTA |
| SAM22-16 | ACGTGAGAGCAAGTCCGTGGTAGGGCAGGTTGGGGTGACT ACATACGACGA | [FAM]-TCTCACGTTCG TCGTA |
| SAM20-18 | ACGTGAGAGCAGTCCGTGGTAGGGCAGGTTGGGGTGACT CATACGACGA | [FAM]-CTCTCACGTTCG TCGTAT |
| SAM22-18 | ACGTGAGAGCAAGTCCGTGGTAGGGCAGGTTGGGGTGACT ACATACGACGA | [FAM]-CTCTCACGTTCG TCGTAT |
| SAM22-20 | ACGTGAGAGCAAGTCCGTGGTAGGGCAGGTTGGGGTGACT ACATACGACGA | [FAM]-GCTCTCACGT TCGTCGTATG |
| SAM16 with gap = 0 | ACGTGAGAAGTCCGTGGTAGGGCAGGTTGGGGTGACT TACGACGA | [FAM]-TCTCACGT TCGTCGTA |
| SAM16 with gap = 1 | ACGTGAGAAGTCCGTGGTAGGGCAGGTTGGGGTGACT TACGACGA | [FAM]-TCTCACGT T TCGTCGTA |
| SAM16 with gap = 2 | ACGTGAGAAGTCCGTGGTAGGGCAGGTTGGGGTGACT TACGACGA | [FAM]-TCTCACGT TT TCGTCGTA |
| SAM16 with gap = 3 | ACGTGAGAAGTCCGTGGTAGGGCAGGTTGGGGTGACT TACGACGA | [FAM]-TCTCACGT TTT TCGTCGTA |
| SAM16-16 with EcoRI site | [FAM]-AATTCCGA AGTCCGTGGTAGGGCAGGTTGGGGTGACT ATCGACGG | TAGCTGCCTTAAGGCT |
